# Individual differences in cortical processing speed predict cognitive abilities: a model-based cognitive neuroscience account

**DOI:** 10.1101/374827

**Authors:** Anna-Lena Schubert, Michael D. Nunez, Dirk Hagemann, Joachim Vandekerckhove

## Abstract

Previous research has shown that individuals with greater cognitive abilities display a greater speed of higher-order cognitive processing. These results suggest that speeded neural information-processing may facilitate evidence accumulation during decision making and memory updating and thus yield advantages in general cognitive abilities. We used a hierarchical Bayesian cognitive modeling approach to test the hypothesis that individual differences in the velocity of evidence accumulation mediate the relationship between neural processing speed and cognitive abilities. We found that a higher neural speed predicted both the velocity of evidence accumulation across behavioral tasks as well as cognitive ability test scores. However, only a small part of the association between neural processing speed and cognitive abilities was mediated by individual differences in the velocity of evidence accumulation. The model demonstrated impressive forecasting abilities by predicting 36% of individual variation in cognitive ability test scores in an entirely new sample solely based on their electrophysiological and behavioral data. Our results suggest that individual differences in neural processing speed might affect a plethora of higher-order cognitive processes, that only in concert explain the large association between neural processing speed and cognitive abilities, instead of the effect being entirely explained by differences in evidence accumulation speeds.

## Introduction

Individual differences in cognitive abilities are important predictors for real-world achievements such as job performance and highest level of educational attainment (Schmidt & Hunter, 2004). Cognitive ability differences also predict differences in individuals’ health (Deary, 2008; Der, Batty, & Deary, 2009), happiness (Nikolaev & McGee, 2016), and well-being (Pesta, McDaniel, & Bertsch, 2010). However, what remains largely unexplored are the fundamental biological processes that give rise to individual differences in cognitive abilities across individuals. In this study we explore how individual differences in cognitive abilities are associated with individual differences in neural processing speed, and how this association can be explained by individual differences in the velocity of evidence accumulation as an intermediate cognitive process.

Previous research has suggested that those individuals with greater cognitive abilities have a higher speed of information-processing, typically measured as reaction or inspection times in elementary cognitive tasks on a behavioral level (Kyllonen & Zu, 2016; Sheppard & Vernon, 2008), or as latencies of event-related potential (ERP) components on a neurophysiological level (e.g., Bazana & Stelmack, 2002; Schubert, Hagemann, Voss, Schankin, & Bergmann, 2015; Troche, Indermühle, Leuthold, & Rammsayer, 2015). Neuroimaging studies have shown that the association between the speed of information-processing and cognitive abilities may reflect individual differences in white-matter tract integrity, either as an overall brain property (Penke et al., 2012) or in specific brain regions such as the forceps minor and the corticospinal tract (Kievit et al., 2016).

However, those with greater cognitive abilities do not seem to benefit from a higher speed of information-processing during *all* stages of information-processing. Instead, individuals with greater cognitive abilities show a higher speed of information processing only in higher-order cognitive processes such as decision making and memory updating (Schmiedek, Oberauer, Wilhelm, Suss, & Wittmann, 2007; Schubert, Hagemann, & Frischkorn, 2017). In particular, the velocity of evidence accumulation during decision making has been repeatedly associated with individual differences in cognitive abilities (Schmiedek et al., 2007; Schmitz & Wilhelm, 2016; Schubert et al., 2015; van Ravenzwaaij, Brown, & Wagenmakers, 2011). Moreover, cognitive abilities have been specifically associated with the latencies of ERP components reflecting higher-order cognitive functions such as memory and context updating (Bazana & Stelmack, 2002; McGarry-Roberts, Stelmack, & Campbell, 1992; Schubert et al., 2017; Troche, Houlihan, Stelmack, & Rammsayer, 2009). Taken together, these results suggest that a greater speed of information-processing may facilitate evidence accumulation during decision making and memory updating and may give rise to advantages in general cognitive abilities. In the present study, we explore this hypothesis by using a hierarchical Bayesian cognitive modeling approach to investigate if individual differences in the velocity of evidence accumulation mediate the relationship between neural processing speed and general cognitive abilities.

### Measuring the speed of higher-order cognitive processes

Reaction time measures are affected by a variety of cognitive and motivational processes and differences across individuals are not solely due to differences in the specific processes of interest (Nunez, Srinivasan, & Vandekerckhove, 2015; Schubert et al., 2015). Therefore, mean reaction times and differences in reaction times between certain experimental conditions can only provide very imprecise measurements of the speed of *specific* specific higher-order cognitive processes. One approach to measure the speed of higher-order cognitive processes is to use validated mathematical models of decision making, which allow estimating the speed and efficiency of specific cognitive processes (Voss, Rothermund, &Voss, 2004). One of the most influential model types used to jointly describe reaction time distributions and accuracies in binary choice tasks are diffusion models. Diffusion models assume that information accumulation follows a continuous, stochastic Wiener process that terminates once one of two decision thresholds has been reached (Ratcliff, 1978; Ratcliff & McKoon, 2008; Stone, 1960). That is, it is assumed that on any given trial an individual will accumulate evidence for one choice over another in a random walk evidence accumulation process with an infinitesimal time step (while neural coding may be more sequential in nature, the infinitesimal approximation should hold true for small time steps). It is predicted that the change in relative evidence *E_t_* follows a Wiener (i.e., Brownian motion) process with an average evidence accumulation rate *δ* and instantaneous variance *ς*^2^ (Ross, 2014) such that

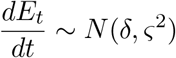

Typically, the variance *ς*^2^ is fixed to some standardized value for reasons of identifiability (but see Nunez, Vandekerckhove, & Srinivasan, 2017). The drift rate (*δ*) measures the relative velocity of evidence accumulation during decision making and individual differences in this parameter have been suggested to be associated with individual differences in cognitive abilities (Ratcliff, Thapar, & McKoon, 2010, 2011; Schmiedek et al., 2007; Schmitz & Wilhelm, 2016; Schubert et al., 2015). The evidence units per second of the drift rate (*δ*) are relative to a predetermined decision criterion for evidence (*α*), which reflects speed-accuracy trade-offs (Voss et al., 2004). In addition, a basic diffusion model consists of one more additional parameter describing and complementing the decision process: The non-decision time (*t_er_*) encompasses all non-decisional processes such as encoding and motor reaction time.

It is not surprising that the drift rate parameter in particular has become widely popular in individual differences research, because it allows quantifying the speed of information uptake free of confounding process parameters such as encoding and motor times or decision cautiousness, which are captured by other model parameters and are largely irrelevant for cognitive abilities research. Individual differences in drift rates have been shown to exhibit trait-like properties (i.e., they show temporal stability and trans-situational consistency; Schubert, Frischkorn, Hagemann, & Voss, 2016) and to be associated with individual differences in cognitive abilities (Ratcliff et al., 2010, 2011; Schmiedek et al., 2007; Schmitz & Wilhelm, 2016; Schubert et al., 2015), attention (Nunez et al., 2015), and word recognition (Yap, Balota, Sibley, & Ratcliff, 2012). The drift rate can even be interpreted in the framework of item response theory (IRT), in which it can under certain assumptions be decomposed into an ability and difficulty parameter (van der Maas, Molenaar, Maris, Kievit, & Borsboom, 2011).

Moreover, several studies suggest a direct link between drift rates and neural processing correlates in the EEG. In particular, it has been shown that the P3, an ERP component occurring typically about 250-500 ms after stimulus onset with a positive deflection that is maximal at parietal electrodes (Polich, 2007), is a neural correlate of the evidence accumulation process captured in the drift rate (Kelly & O’Connell, 2013; O’Connell, Dockree, & Kelly, 2012; Ratcliff, Philiastides, & Sajda, 2009; Ratcliff, Sederberg, Smith, & Childers, 2016; van Ravenzwaaij, Provost, & Brown, 2017). O’Connell et al. (2012) and Kelly and O’Connell (2013) even suggested that the buildup rate of this positive centroparietral positive potential may directly reflect the rate of evidence accumulation on a neural level.

Particularly intriguing from an individual-differences perspective is the observation that individual differences in P3 amplitudes across conditions have been shown to explain about 74 percent of the variance in drift rates *δ* (Ratcliff et al., 2009). Because both individual differences in drift rates and individual differences in P3 characteristics have been shown to explain cognitive abilities, it may be proposed that the relationship between ERP latencies reflecting higher-order cognition and cognitive abilities is mediated by individual differences in drift rates. In particular, we hypothesize that a greater speed of neural information processing positively affects evidence accumulation, and that this facilitation of evidence accumulation translates to advantages in general cognitive abilities. While it has been previously shown that reaction times partly mediate the relationship between ERP latencies and cognitive abilities (Schubert et al., 2015), no studies have yet jointly analyzed individual differences in *evidence accumulation rates*, ERP latencies, and cognitive abilities. Instead, mathematical modeling and cognitive neuroscience have independently contributed to our understanding of the basic processes underlying individual differences in cognitive abilities. While mathematical modeling has suggested that the velocity of evidence accumulation may be specifically related to cognitive abilities (Ratcliff et al., 2010, 2011;Schmiedek et al., 2007; Schubert et al., 2015), cognitive neuroscience approaches have characterized the time-course of information-processing and identified structural and function neural correlates of cognitive abilities (Basten, Hilger, & Fiebach, 2015; Jung & Haier, 2007; Neubauer & Fink, 2009). Recent advancements in the emerging field of model-based cognitive neuroscience have demonstrated the advantages of integrating mathematical modeling and cognitive neuroscience by contributing to a deeper theoretical understanding of multiple modes of data and allowing better explanation and prediction of new data (e.g., Forstmann, Wagenmakers, Eichele, Brown, & Serences, 2011; Nunez et al., 2017; Palmeri, Love, &Turner, 2017; Turner, Forstmann, Love, Palmeri, & van Maanen, 2017). In this study we use model-based cognitive neuroscience techniques to find evidence for or against our hypothesis.

### A model-based cognitive neuroscience account of individual differences in cognitive abilities

Jointly analyzing behavioral and brain data improves inferences about human cognition, because it is assumed that both measures reflect properties of the same latent cognitive process. This simultaneous analysis can be achieved in a hierarchical Bayesian framework using formal mathematical models such as the diffusion model to constrain or inform inferences based on the brain data (Forstmann et al., 2011; Turner et al., 2017). The hierarchical Bayesian framework provides many advantages (M. D. Lee, 2011; Shiffrin, Lee, Kim, & Wagenmakers, 2008). First and foremost, joint models are fit to all data simultaneously and do not require separate parameter estimation stages that lead to an underestimation of parameter uncertainty or standard errors (Vandekerckhove, 2014). Both empirical and simulation studies have shown that ignoring the hierarchy in hierarchically structured data can bias inferences drawn from these data (Boehm, Marsman, Matzke, & Wagenmakers, 2018; Vandekerckhove, 2014)

Second, hierarchical Bayesian models can easily handle low observation counts or missing data structures (M. D. Lee & Wagenmakers, 2014), which is an ideal property when the cost of collecting neural measurements is high. In particular, Bayesian Markov Chain Monte Carlo (MCMC) sampling finds posterior distributions of model parameters without the need for strong assumptions regarding the sampling distribution of these parameters (Levy & Choi, 2013). Moreover, Bayesian statistical modeling approaches do not rely on asymptotic theory (S. Y. Lee & Song, 2004). These two properties make convergence issues in multivariate regression models in smaller samples less likely. Another favorable property of Bayesian hierarchical modeling is *shrinkage*, which describes the phenomenon that individual parameter estimates are informed by parameter estimates for the rest of the sample. Because less reliable and outlier estimates are pulled towards the group mean, shrinkage has been used in neuroimaging research to improve the reliability of individual functional connectivity estimates by 25 to 30 percent (Dai & Guo, 2017; Mejia et al., 2018; Shou et al., 2014). Taken together, these desirable properties of hierarchical Bayesian models open up the possibility to use multivariate regression models such as structural equation models (SEM) or latent growth curve models in neuroimaging research, where sample sizes are usually smaller than in behavioral research due to the cost associated with the collection of neural measures.

The joint analysis of behavioral and neural data can be expanded into a cognitive latent variable model (CLVM) by including data from multiple conditions and/or tasks and by introducing covariates such as cognitive ability tests or personality questionnaires into the hierarchical model (Vandekerckhove, 2014; Vandekerckhove, Tuerlinckx, & Lee, 2011). In addition to jointly modeling behavioral and neural data, the cognitive latent variable framework allows estimating correlations between higher-order variables, which reflect the covariances between behavioral, neural, and cognitive abilities data across experimental tasks or ability tests. As such, a CLVM is a computationally expensive, but highly flexible tool that strongly resembles structural equation modeling (SEM) in the way that it allows specifying associations between latent variables and distinguishing between constructs and their measurements. Vandekerckhove (2014) demonstrated the advantages of a CLVM in comparison to a more conventional two-stage analysis when modeling the latent association between evidence accumulation rates in executive function tasks and psychometric measures of dysphoria.

In the present study, we constructed CLVMs to assess the latent relationship between latencies of ERP components reflecting higher-order processing (P2, N2, P3), reaction times and accuracies in elementary cognitive tasks, and general cognitive abilities (see **Figure 1**). For this purpose, we reanalyzed data from a study with multiple measurement occasions previously reported in Schubert et al. (2017). In particular, we wanted to test if the association between latencies of ERP components associated with higher-order cognitive functions and general cognitive abilities established with conventional structural equation modeling could be explained by individual differences in the velocity of evidence accumulation. We also conducted out-of-sample forecasts to validate how well this mediation model was able to predict individual cognitive ability test scores solely based on new participants electrophysiological and behavioral data. We expected that a greater speed of neural information-processing would facilitate evidence acquisition during decision making and memory updating, and that this advantage in the velocity of evidence accumulation would mediate the predicted association between neural processing speed and general cognitive abilities.

**Figure 1.**
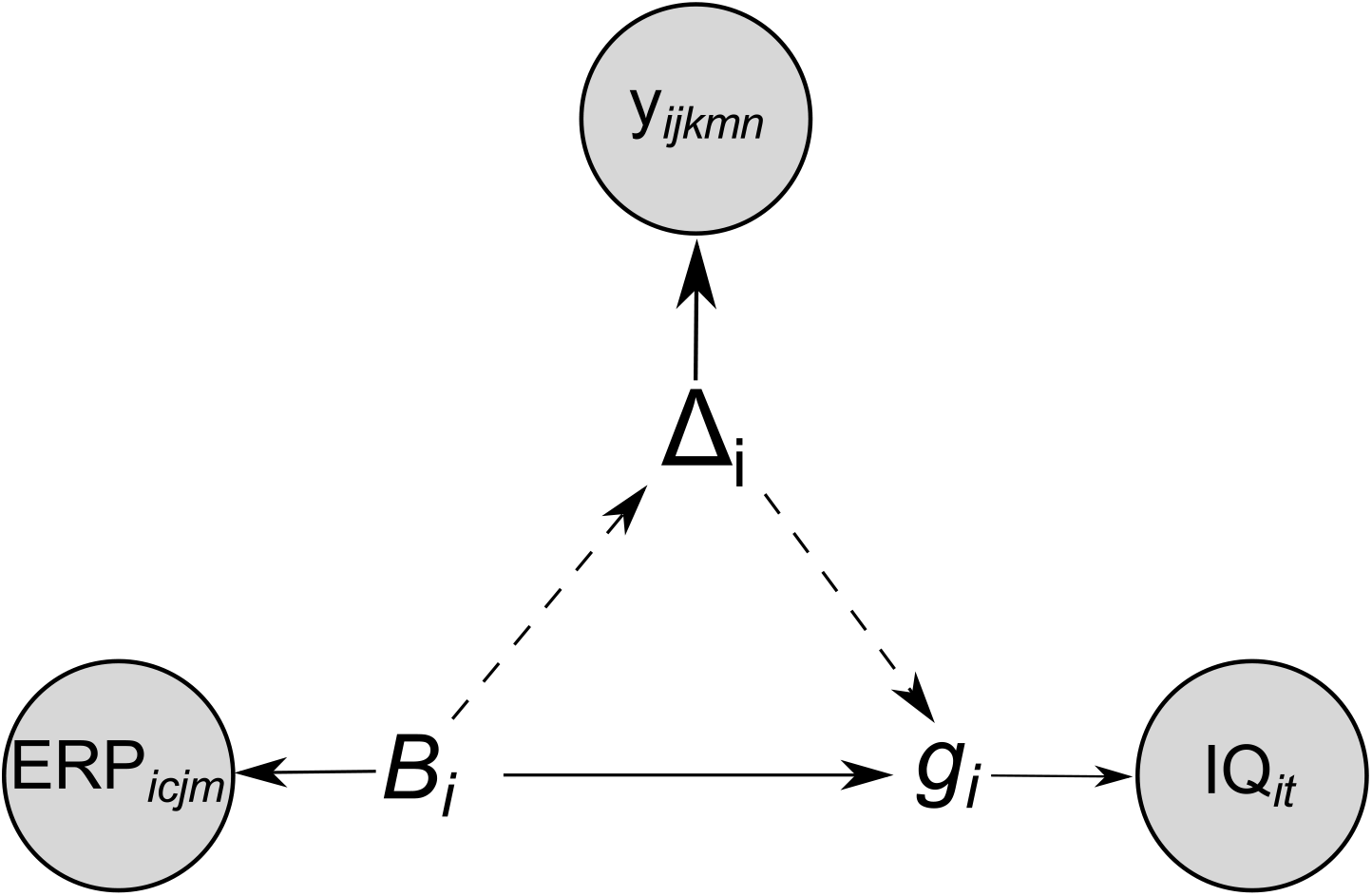
Simple visualization of both linking models (such that the mediation-linking model includes dashed connections). Shaded nodes represent observed data across participants i. *B_i_*, *δ_I_*, and *g_i_* represent the highest latent variables of neural processing speed (left: describing shared variance across ERP latencies), evidence accumulation velocity (top: describing shared variance across reaction time distributions), and cognitive ability (right: describing shared variance across intelligence test scores).

## Method

### Participants

*N* = 122 participants (72 females, 50 males) from different occupational and educational backgrounds participated in three sessions of the study. They were recruited via local newspaper advertisements, social media platforms, and flyer distributions in the Rhine-Neckar metropolitan region. Participants were between 18 and 60 years old (*M* = 36.7, *Med* = 35.0, *SD* = 13.6), had normal or corrected to normal vision, and reported no history of mental illness. All participants signed an informed consent prior to their participation in the experiment. The study was approved by the ethics committee of the faculty of behavioral and cultural studies, Heidelberg University.

### Procedure

The study consisted of three sessions that were each approximately four months apart. Participants completed the experimental tasks in the first and third session while their EEG was recorded in a dimly-lit, sound-attenuated cabin. The order of tasks (choice reaction time task, recognition memory task, letter matching task) was the same for all participants and both sessions. During the second session, participants completed the cognitive ability tests, a personality questionnaire (Kretzschmar, Spengler, Schubert, Steinmayr, & Ziegler, 2018), and a demographic questionnaire. Each session lasted approximately 3-3.5 hours in duration with EEG being collected for approximately 2.5 hours. Participants were given breaks between tasks and conditions to reduce mental fatigue.

### Measures

#### Experimental tasks

##### Choice reaction time task (CR)

Participants completed a choice reaction time task with two conditions, a two-alternative (CR2) and a four-alternative (CR4) choice condition. Four white squares were presented in a row on a black screen. Participants’ middle and index fingers rested on four keys directly underneath the squares. After a delay of 1000-1500 ms, a cross appeared in one of the four squares and participants had to press the corresponding key as fast and accurate as possible. The screen remained unchanged for 1000 ms after their response to allow the recording of post-decision neural processes. Then, a black screen was shown for 1000-1500 ms between subsequent trials; the length of the inter-trial interval (ITI) was uniformly distributed. See the left part of **Figure 2** for an overview of the experimental procedure.

**Figure 2.**
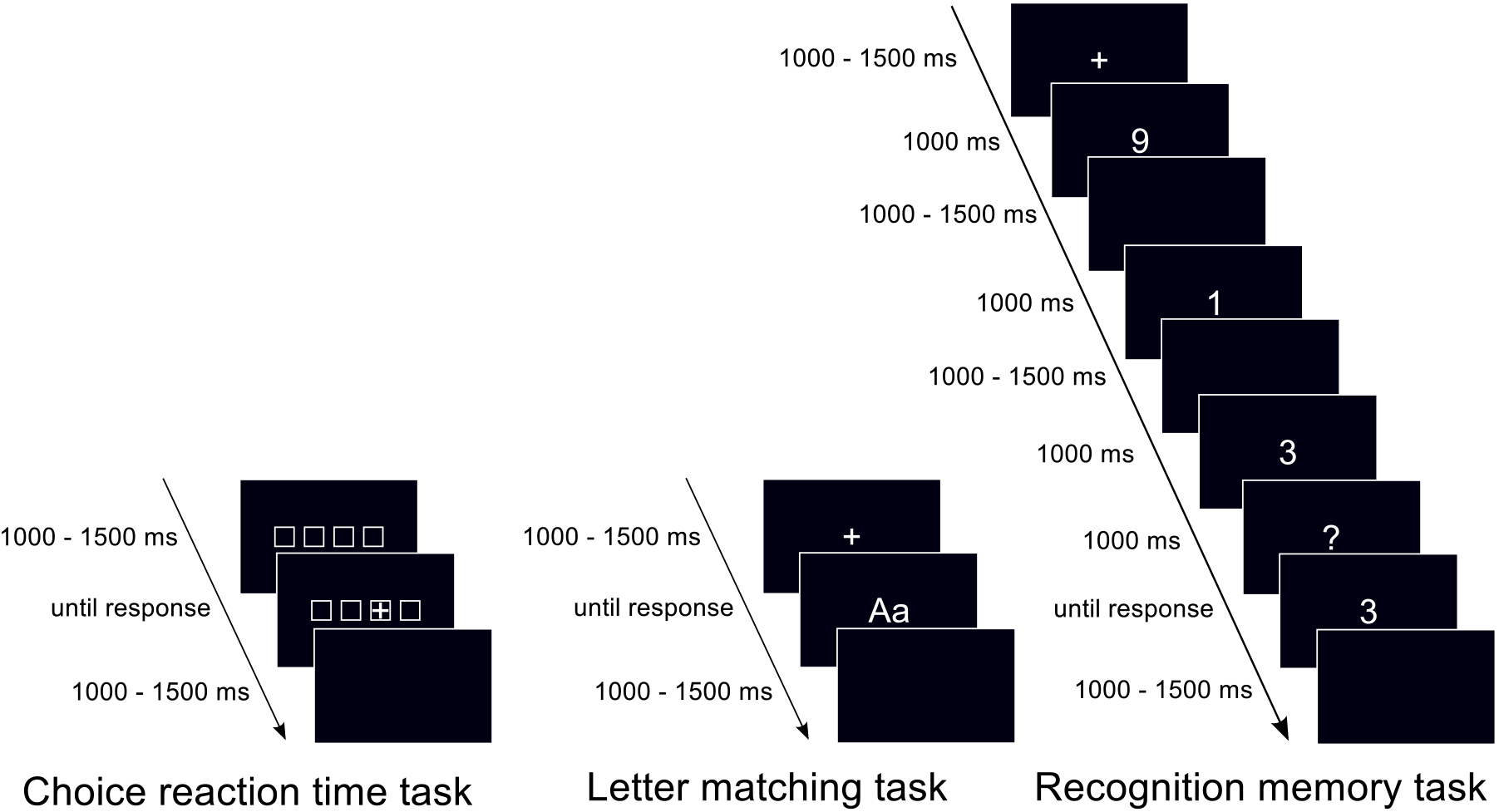
Participants completed three experimental tasks. The choice reaction time task (CR) consisted of a 2-choice (CR2) and a 4-choice (CR4) condition with 200 trials each, the recognition memory task of a physical identity (PI) and name identity (NI) condition with 300 trials each, and the recognition memory task (RM) of memory set sizes 1 (RM1), 3 (RM3), and 5 (RM5) with 100 trials each.

In the two-choice response time condition, the number of choices was reduced to two squares in which the cross could appear for 50 subsequent trials. In the four-choice response time condition, the cross could appear in any of the four squares. Both conditions began with ten practice trials with immediate feedback followed by 200 test trials without feedback. The order of conditions was counterbalanced across participants.

##### Letter matching task (LM)

Participants saw two white letters on a black screen and had to decide whether they were physically (physical identity condition) or semantically (name identity condition) identical by pressing one of two keys. Letters were identical in 50% of the trials. Each trial was followed by an inter-trial interval (ITI) of 1000-1500 ms. See the middle part of **Figure 2** for an overview of the experimental procedure. Conditions were presented block-wise. Each condition began with ten practice trials with immediate feedback followed by 300 test trials without feedback. All participants completed the physical identity condition first at the first measurement occasion, and second at the second measurement occasion.

##### Recognition memory task (RM)

Participants viewed memory sets of white, numerical digits (0 to 9) on a black screen. Digits were presented sequentially for 1000 ms each followed by a blank inter-stimulus interval shown for 400-600 ms. After the final digit was presented, participants saw a black screen with a white question mark for 1800-2200 ms. Subsequently, they were shown a single digit and had to decide whether the digit had been included in the previously presented memory set by pressing one of two keys. Each trial was followed by a uniformly distributed ITI of 1000-1500 ms. The probe digit was included in the memory set in 50% of the trials. There were three conditions of the experiment with the memory set consisting of either one, three, or five digits. See the right part of **Figure 2** for an overview of the experimental procedure in the set size 3 condition. The three conditions were presented block-wise and the order of presentation was counterbalanced across participants. Each condition consisted of ten practice trials with immediate feedback followed by 100 test trials without feedback.

#### Cognitive abilities tests

##### Berlin intelligence structure test (BIS)

We administered the Berlin intelligence structure test (Jäger & Süß, 1997), which distinguishes between four operation-related (processing speed, memory, creativity, processing capacity), and three content-related (verbal, numerical, figural) components of cognitive abilities. Each of the 45 tasks included in the test consists of a combination of one operation-with one content-related component. Following the manual, we calculated participants’ scores in the four operation-related components by aggregating the normalized *z*-scores of tasks reflecting the specific operational components irrespective of content. The mean score of the processing capacity (PC) component was *M* = 101.70 (*SD* = 7.99), the mean score of the processing speed (PS) component was *M* = 98.00 (*SD* = 7.10), the mean score of the memory (M) component was *M* = 99.40 (*SD* = 6.51), and the mean score of the creativity (C) component was *M* = 98.02 (*SD* = 6.14). We then transformed these scores to *z*-scores for further analyses.

##### Advanced Progressive Matrices (APM)

Participants completed a computer-adapted version of Raven’s Advanced Progressive Matrices (Raven, Court, & Raven, 1994). The APM is a fluid intelligence test that consists of 36 items. Each item consists of a 3×3-matrix with geometric figures that follow certain logical rules and symmetries. The last element of the matrix is missing and must be chosen out of eight alternatives without time limit (see **Figure 3** for a fictional sample item). Following the manual, participants’ performance was calculated as the number of correctly solved items of the second set. Moreover, we calculated performance in the odd and even trials of the test separately to construct two indicators of latent APM performance. We then transformed these raw test sores to *z*-scores for further analyses. Participants solved on average *M* = 23.43 (*SD* = 6.71) items correctly, which corresponds to a mean IQ score of *M* = 98.80 (*SD* = 15.68). Performance on even trials, *M_even_* = 12.23 (*SD* = 3.51) correctly solved items, was comparable to performance on odd trials, *M_odd_* = 11.20 (*SD* = 3.52) correctly solved items.

**Figure 3.**
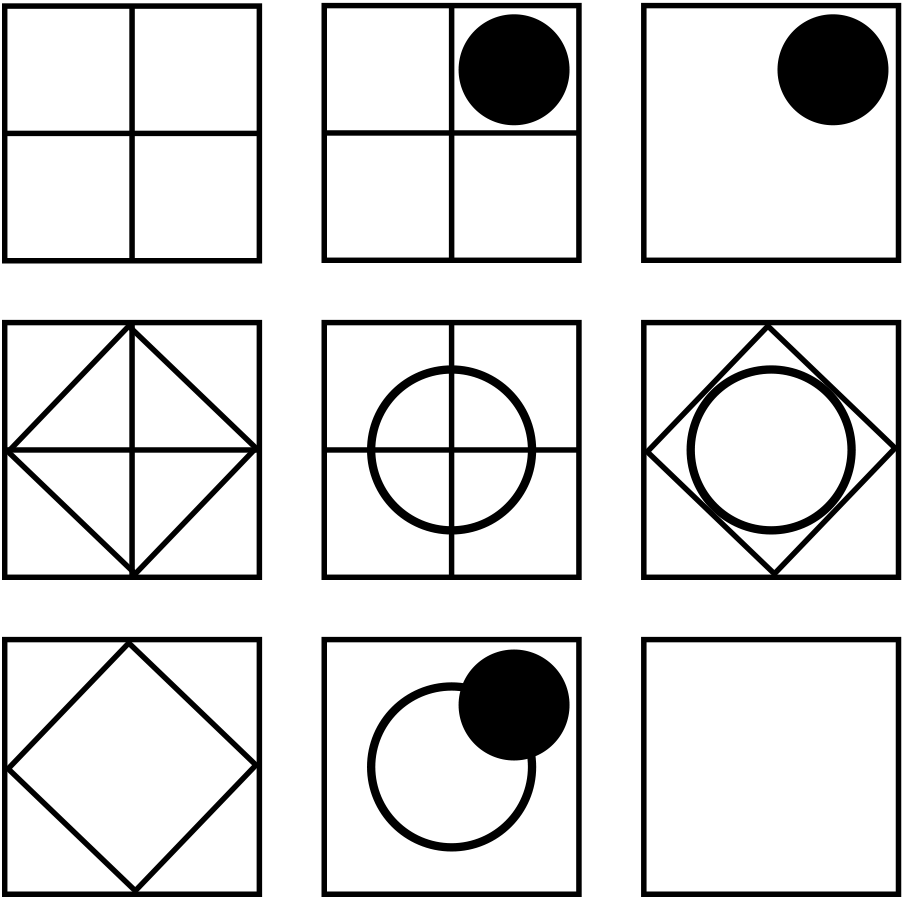
Example stimuli of Raven’s Progressive Matrices. Each item consists of a 3×3-matrix with geometric figures that follow certain logical rules and symmetries. The last element of the matrix is missing and must be chosen out of eight alternatives.

### EEG recording

Participants’ EEG was recorded with 32 equidistant silver-silver chloride electrodes, a 32-channel BrainAmp DC amplifier (Brain Products, Munich) and a sampling rate of 1000 Hz (software bandpass filter of 0.1-100 Hz with a slope of 12 db/octave). In addition, participants’ electrooculogram (EOG) was recorded bipolarly with two electrodes positioned above and below the left eye and two electrodes positioned at the outer corners of the eyes. Electrode impedances were kept below 5 kΩ during recording. Data were collected with a central electrode reference but later offline re-referenced to the average activity of all electrodes (average reference). The data were filtered offline with a low-pass filter of 16 Hz with a slope of 12 db/octave.

### Data analysis

#### Behavioral data

To remove outliers in the behavioral data, we discarded any reaction times faster than 100 ms or slower than 3000 ms. In a second step, we discarded any trials with logarithmized reaction times exceeding ± 3 standard deviations from the mean reaction time of each condition. Deviations in criteria (i.e., less strict criteria) did not affect the covariance structure between variables, suggesting adequate robustness.

#### Evoked electrophysiological measures

Event-related potentials (ERPs) were analyzed separately for each task and condition. ERPs were calculated by averaging all experimental trials, time-locked to the onset of the task-relevant visual stimuli, with windows of interest that were 1000 ms long with a preceding baseline of 200 ms. Ocular artifacts were corrected for with the regression procedure suggested by Gratton, Coles, and Donchin (1983). Windows of EEG data with amplitudes exceeding ± 70 *μV* at least once within the time window, with amplitude changes exceeding 100 *μV* within 100 ms, or with activity lower than 0.5 *μV* were discarded as artifacts.

Latencies of three ERP components were calculated for each participant in each experiment. Grand-average waveforms of event-related potentials are presented in **Figure 4**. P2 peak latencies were determined with regard to the greatest positive local maxima at the fronto-central electrode on the midline, which roughly corresponds to the Fz electrode in the 10-20 system, in a 120 to 320 ms time window. N2 and P3 peak latencies were determined with regard to the greatest negative and positive local maxima at the parietal electrode on the midline, which roughly corresponds to the Pz electrode in the 10-20 system, in a 140 to 370 ms time window (N2) and a 200 to 630 ms time window (P3), respectively. Peak latencies were determined separately for each condition of each experimental task, then averaged across conditions within each experiment, and then *z*-standardized for further analyses. Prior to averaging across experimental conditions, we discarded any peak latencies exceeding ± 3 SDs from the mean peak latency of each condition. If any peak latencies were discarded, the average across conditions was calculated based on the remaining conditions.

**Figure 4.**
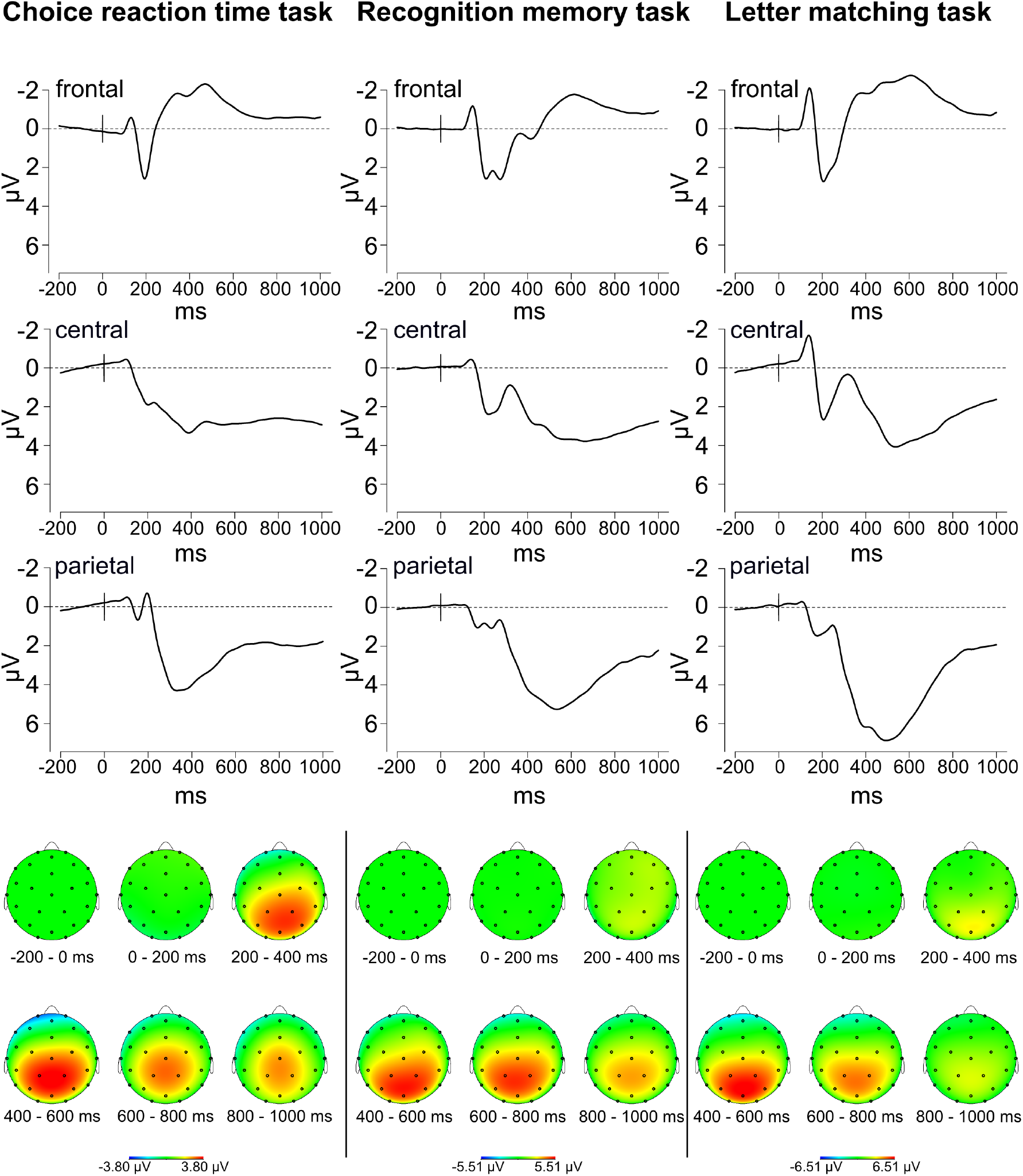
Grand averages of event-related potentials at frontal, central, and parietal electrodes over midline. ERPs were elicited by stimulus onset and averaged across laboratory sessions and conditions for each experimental task.

#### Cognitive latent variable models

We constructed hierarchical Bayesian models to assess the latent relationship between reaction times, latencies of the three ERP components (P2, N2, P3), and cognitive ability test scores. For this purpose, we defined three separate sub-models describing the domain-specific associations between a) ERP latencies in experimental tasks across two measurement occasions, b) behavioral data in experimental tasks across two measurement occasions, and c) performance in cognitive ability tests.

Then, we constructed two models using either 1) only ERP latencies or 2) ERP latencies and behavioral data to predict performance in cognitive ability tests. To test the hypothesis that drift rates mediate the relationship between neural processing speed and cognitive abilities, we compared performance of a direct regression model, in which ERP latencies predicted cognitive abilities (“Direct Regression Model”), to a mediation model, in which the effect of ERP latencies on cognitive abilities was mediated by drift rates (“Mediation Model”).

We used Just Another Gibbs Sampler (JAGS; Plummer, 2003) with a module that adds a diffusion model distribution to JAGS (jags-wiener; Wabersich & Vandekerckhove, 2014) to find parameter estimates for the hierarchical model. Each model was fit with three Markov Chain Monte Carlo (MCMC) chains run in parallel. Each chain contained 2,000 burn-in samples and 100,000 additional samples with a thinning parameter of 10, resulting in 10,000 posterior samples per chain. Posterior samples from the three chains were combined to one posterior sample consisting of 30,000 samples for each model parameter. Model convergence was evaluated based on the Gelman-Rubin convergence statistic R̂, which compares the estimated between-chains and within-chain variances for each model parameter (Gelman & Rubin, 1992). Negligible differences between these variances were indicated by R̂ values close to 1.

##### Submodel: ERP latencies in experimental tasks

ERP latencies were modeled in a hierarchical structural equation model (SEM) inspired by the parameter expansion approach suggested by Merkle and Rosseel (2018). Each of the three ERP latencies (P2, N2, P3) was quantified in three tasks at two sessions. Hence, six observed variables (3 tasks *j* × 2 sessions *m*) loaded onto each one of the three the first-order component *c* specific ERP factors *η*_(*P*2)_, *η*_(*N*2)_, and *η*_(*P*3)_. These three latent components loaded onto a second-order latent factor *B* that was estimated per participant *i*.

Latent factors and observed variables had normally distributed prior and hyperprior distributions. The means of these priors reflected linear regressions of the respective higher-order factors. For reasons of identifiability, the loading *γ*_(*P*2)_ of the first lower-order factor *η_P_*_2_ on the higher-order factor *B* was fixed to 1, while the other loadings, *γ*_(*N*2)_ and *γ*_(*P*3)_, were given standard normal priors: *γ*_(*P*2)_ = 1 and *γ*_(*N*2)_*, γ*_(*P*3)_ ~ *𝒩* (0, 1).

Finally, precisions *ψ* (inverses of variances) of all latent variables were modeled as gamma distributed variables: *Ψ*_*B*_, *ψ*_(*P*2)_, *ψ*_(*N*2)_, *ψ*_(*P*3)_ ~ Γ(1, 0.5).

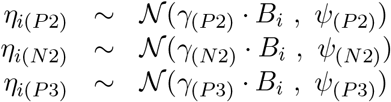

For the second-order latent factor,

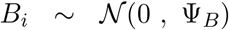

Subsequently, the observed latencies *ERP_icjm_* of ERP components *c*, tasks *j*, and measurement occasions *m* for each participant *i* were regressed onto the first-order latent variables. These regressions were defined by the respective factor loadings *λ_cjm_*, the respective higher-order latent variables *η_ic_*, and the respective precisions *θ_cjm_*. Factor loadings *λ_cjm_* on the first-order latent variables were fixed to 1 for task *j* = *CR* and measurement occasion *m* = 1 for all three ERP components for reasons of identifiability. See the bottom left parts of **Figure 5**, **Figure 6**, and **Figure 7** for a graphical illustration of the measurement model of ERP latencies.

**Figure 5.**
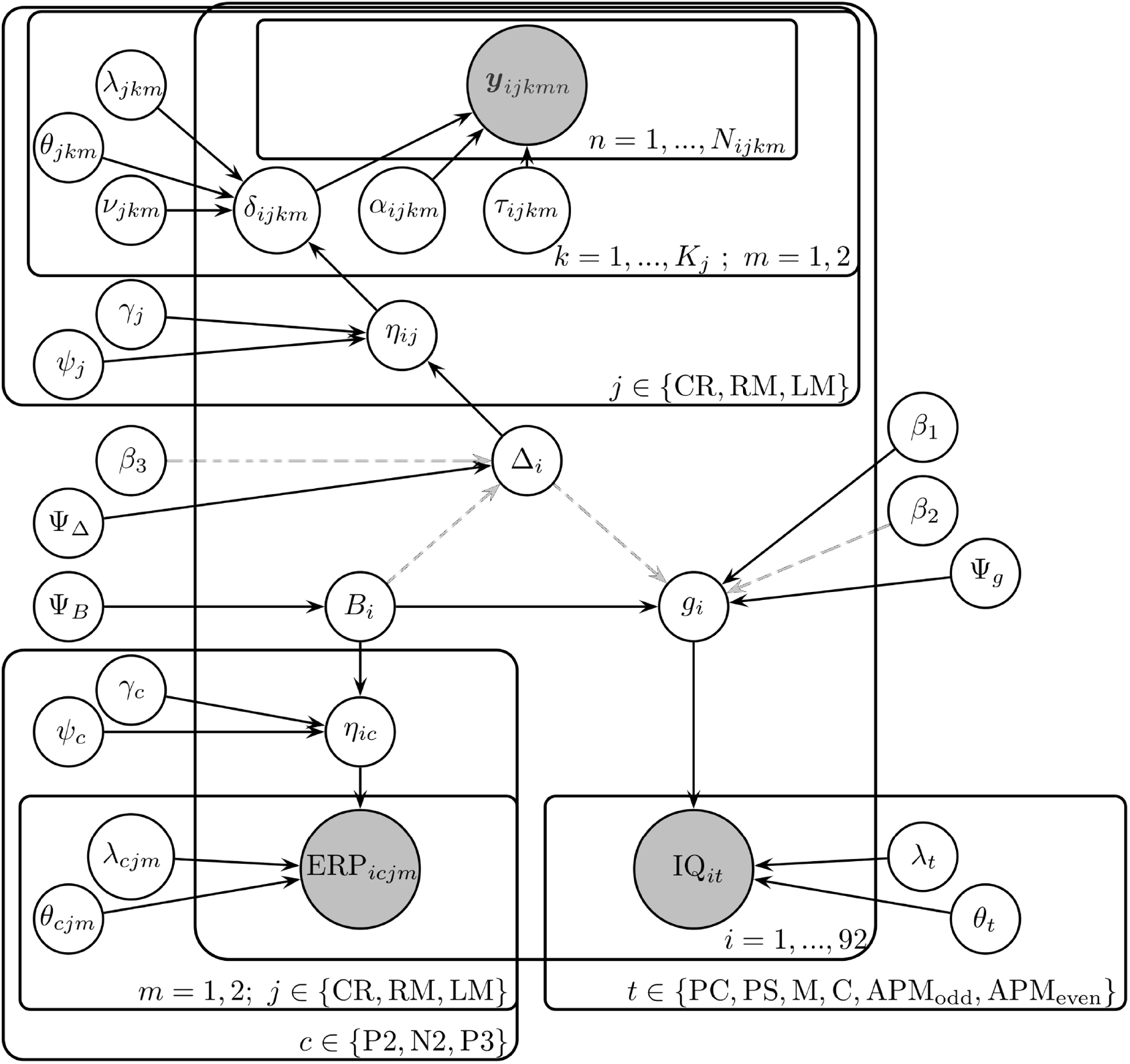
Graphical visualization of both the regression-linking and mediation-linking models (such that the mediation-linking model includes dashed connections). An alternate way of understanding the neurocognitive models presented in this manuscript is by viewing the graphical notation for hierarchical models as described by M. D. Lee and Wagenmakers (2014). Shaded notes represent observed data while unshaded nodes represent unknown (fitted) parameters. Arrows represent direction of influence such that hierarchical parameters influence lower level parameters and observed data. Plates denote the number of observations for each variable and data point of participant *i*, experimental task *j*, experimental condition *k*, measurement occasion *m*, ERP component *c*, cognitive abilities task *t*, and trial *n*. Behavioral data *y* is colored red to indicate that it is a vector of both reaction time and accuracy observations.

**Figure 6.**
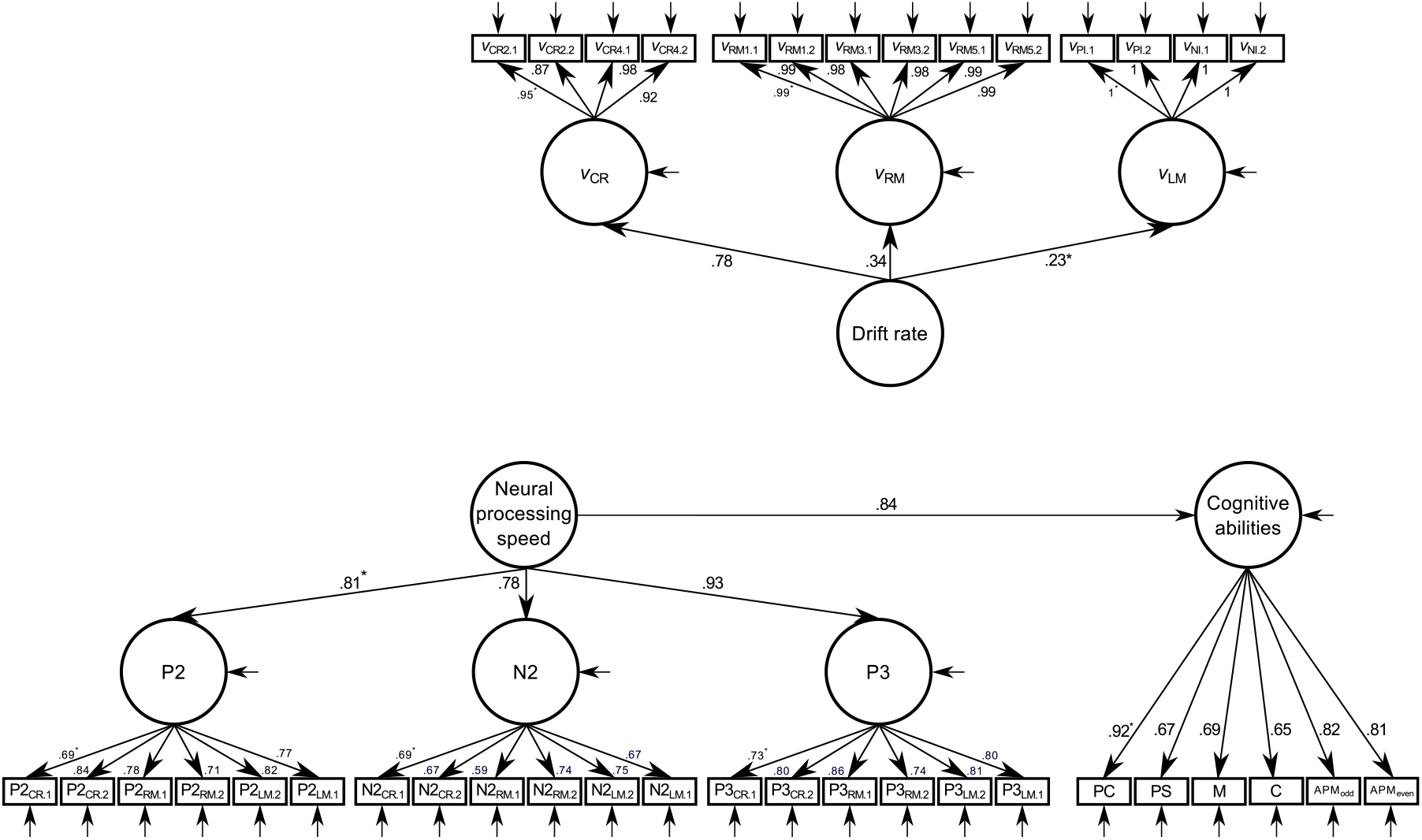
Structural equation modeling visualization of the regression linking model. Posterior medians of standardized regression weights are shown next to paths. Asterisks indicate factor loadings fixed to 1. CR/CR2/CR4 = choice reaction time task with two or four alternatives; RM/RM1/RM3/RM5 = recognition memory task with memory set size of 1, 3, or 5; LM/PI/NI = letter matching task with physical identity or name identity condition; PC = processing capacity; PS = processing speed; M = memory; C = creativity.

**Figure 7.**
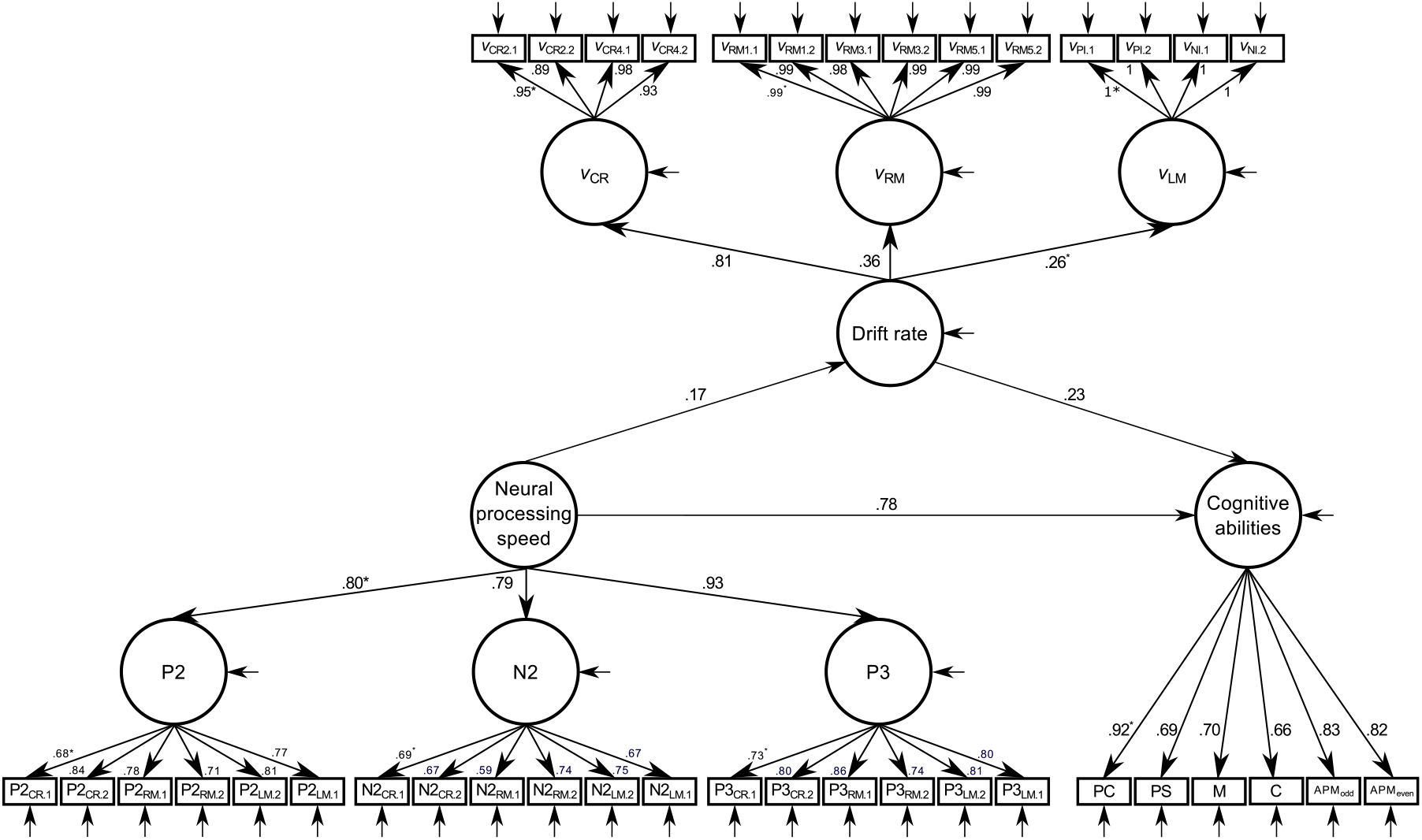
Structural equation modeling visualization of the mediation linking model. Posterior medians of standardized regression weights are shown next to paths. Asterisks indicate factor loadings fixed to 1. CR/CR2/CR4 = choice reaction time task with two or four alternatives; RM/RM1/RM3/RM5 = recognition memory task with memory set size of 1, 3, or 5; LM/PI/NI = letter matching task with physical identity or name identity condition; PC = processing capacity; PS = processing speed; M = memory; C = creativity.

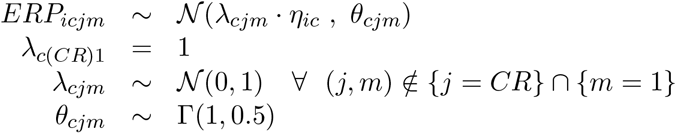

##### Submodel: Behavioral data in experimental tasks

We used a combination of the SEM approach based on parameter expansion described above and the hierarchical diffusion model approach described by Vandekerckhove et al. (2011) to model individual differences in reaction times and accuracies in experimental tasks *j*, conditions *k*, and measurement occasions *m*.

In a first step, we modeled task-, condition-, and measurement occasion-specific drift rates in a hierarchical SEM with three task-specific first-order factors *η_ij_*. These three latent components loaded onto a second-order latent factor ∆_*i*_. Again, latent factors and observed variables had normally distributed priors and hyperpriors. The means of these priors reflected linear regressions of the respective higher-order factors.

For reasons of identifiability, the loading *γ*_(*CR*)_ of the first lower-order factor *η*_(*CR*)_ on the higher-order factor ∆ was fixed to 1, while the other loadings, *γ*_(*RM*)_ and *γ*_(*LM*)_, were given standard normal priors: *γ*_(*CR*)_ = 1 and *γ*_(*RM*)_*, γ*_(*LM*)_ *~ 𝒩* (0, 1). Precisions *ψ* (inverses of variances) of all latent variables were modeled as gamma distributed variables: *ψ*_(*CR*)_, *ψ*_(*RM*)_, *ψ*_(*LM*)_ ~ Γ(1, 0.5).

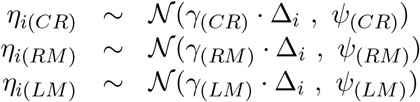

Subsequently, the condition, task-, and measurement-occasion-specific drift rates *δ_ijkm_* were regressed onto the first-order latent variables *η_ij_*. Factor loadings on the respective first-order latent variables were fixed to 1 for condition *k* = 1, referring to the condition with lowest-information processing demands within each task, and measurement occasion *m* = 1 for all three tasks for reasons of identifiability. The other loadings *λ_jkm_* were given standard normal priors: *λ_jkm_* ~ *𝒩* (0, 1). Precisions of drift rates were modeled as gamma distributed variables: *θ_jkm_ ~* Γ(1, 0.5). In addition, we estimated intercepts *ν_jkm_* for the lowest-order drift rates, because the behavioral data were not *z*-standardized: *ν_jkm_* ~ *𝒩* (2, 1.5^2^).

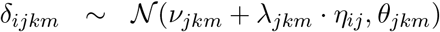

In a second step, these drift rates were entered into the diffusion model distribution in addition to task-, condition-, measurement occasion-, and person-specific boundary separation *α_ijkm_* and non-decision time *τ_ijkm_* parameters (with the starting point parameter fixed at 0.5). Both boundary separation parameters and non-decision times were given standard normal priors: *α_ijkm_* ~ *𝒩* (1, 0.5^2^), *τ_ijkm_* ~ *𝒩* (0.3, 0.2^2^). See the top parts of **Figure 5**, **Figure 6**, and **Figure 7** for a graphical illustration of the measurement model of behavioral data in experimental tasks.

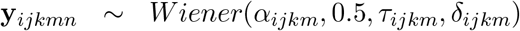

##### Submodel: Performance in cognitive abilities tests

Performance in the two cognitive abilities tests was modeled with a SEM. The four operation-related components of the BIS and the two halves of the APM loaded onto a first-order latent factor *g_i_*.

Subsequently, the observed tests scores *IQ_it_* per cognitive ability test *t* were regressed onto the first-order latent variable *g_i_*. For reasons of identifiability, the loading *λ*_1_ of the processing capacity score of the BIS *η*_1_ on the higher-order factor *g* was fixed to 1, while the other loadings, *λ*_2_, *λ*_3_, *λ*_4_, *λ*_5_, *λ*_6_ were given standard normal priors: *λ*_1_ = 1 and *λ*_2_*, λ*_3_*, λ*_4_*, λ*_5_*, λ*_6_ (0, 1). Precisions *θ* (inverse of variances) of observed IQ scores were given gamma distributed priors: *θ_t_ ~* Γ(1, 0.5). See the bottom right parts of **Figure 5**, **Figure 6**, and **Figure 7** for a graphical illustration of the measurement model of cognitive abilities tests.

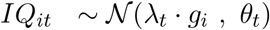

##### Linking models

Finally, we linked all submodels in two linking structures. Whereas the three submodels only established latent *measurement* models for each of the three variable domains (neural data, behavioral data, and cognitive abilities data), the two linking structures specified *structural* associations between variable domains. Hence, the comparison of the two linking models contained the critical comparison: If the velocity of evidence accumulation mediated the relationship between neural speed and cognitive abilities, the mediation model should outperform a direct regression of cognitive abilities on ERP latencies.

We therefore specified two linking structures. In the first linking structure we specified a *regression model* and predicted cognitive abilities tests scores solely through neural processing speed by regressing the latent cognitive abilities factor *g_i_* on the latent ERP latencies factor *B_i_* (see **Figure 1** and compare to **Figure 6**), while the latent drift rate factor ∆_*i*_ was unrelated to the other two latent variables.

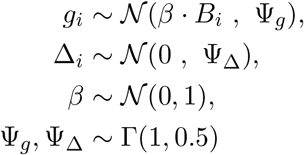

The second linking structure consisted of a *mediation model*, in which the latent cognitive abilities factor *g_i_* was regressed onto both the latent ERP latencies factor *B_i_* and the latent drift rate factor ∆_*i*_, which was in turn regressed onto the latent ERP latencies factor *B_i_* (see **Figure 7**).

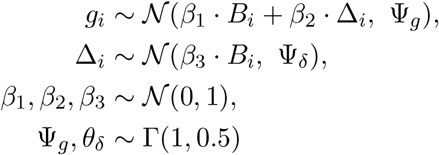

The data of 92 randomly drawn participants (of 114 total; drawn without replacement) were used as a training set to find posterior distributions of cognitive latent variables (i.e., samples from probability distributions that reflect certainty/uncertainty about parameter estimates as reflected by the data). Standardized regression weights were calculated by multiplying unstandardized regression weights with the quotient between the ratio of standard deviation between the predictor (the higher-order latent variable) to the criterion (the lower-order latent or observed variable): 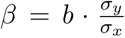. We report the median, 2.5th, and 97.5th percentiles, forming a 95% credible interval (CI) as an equal-tailed interval to describe the posterior distributions of standardized regression weights.

##### Model evaluation

The performance of both linking structures was compared based on their in-sample prediction ability, their Deviance Information Criterion (J., G., P., & Angelika, n.d.), and, crucially, their out-of-sample-prediction ability of new participants data.

*In-sample prediction.* Fitting the model with the training set, we created posterior predictive distributions by simulating new neural, behavioral, and cognitive abilities data separately for each participant based on each participant’s posterior distributions of model parameters and on model specifications. Hence, we simulated two posterior predictive data sets for each of the 92 participants in the training set: One of these posterior predictive data sets was based on model specifications and parameter estimates of the regression model, and the other one based on model specifications and parameter estimates of the mediation model. Subsequently, we assessed how strongly these simulated data were related to the observed data for the whole sample of 92 participants separately for each of the two candidate models. For this purpose, we compared a) observed and predicted ERP latencies for each ERP component *c*, experimental task *j*, and session *m*, b) observed and predicted RT distributions for each condition *c*, experimental task *j*, and session *m*, and c) observed and predicted IQ test scores for each sub-test *t*. RT distributions were compared by comparing the 25th, 50th, and 75th percentile of the observed and predicted RT distributions. To quantify the association between observed and predicted values, we calculated 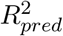 as the proportion of variance of values *T* (ERP latencies, percentiles of the RT distribution, cognitive abilities test scores) explained by model predictions. This statistic is based on the mean squared error of prediction of T, *MSEP_T_*, and the estimated variance of T across participants, 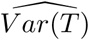.

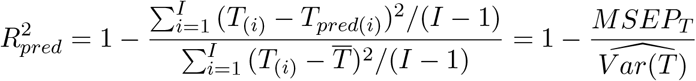

*Deviance information criterion (DIC).* DIC is a measure of goodness-of-fit of a model for hierarchical models that provides a penalty for model complexity (J. et al., n.d.). DIC can be thought of as an extension of Akaike information criterion (AIC) for hierarchical models that enforce shrinkage, such that the number of parameters k is no longer useful as a penalty for model complexity. Another alternative is Bayesian information criterion (BIC), which approximates the logarithm of the Bayes Factor (i.e. the ratio of Bayesian probabilities for two comparison hypotheses), but which is difficult to estimate in most hierarchical models (Kass & Raftery, 1995). Due to ease of estimation and implementation in JAGS (Plummer, 2003), we used DIC as known model comparison metric. Smaller DIC values indicate more favorable models. However, we consider out-of-sample prediction of new participants to be the ultimate test of models that natively penalizes model complexity due to overfitting of in-sample data.

*Out-of-sample prediction.* A test set of 22 new participants (the randomly drawn remaining participants) was used to find a second set of posterior predictive distributions for each participant. This test set allowed us to assess how well models were able to predict new participants’ data in one domain (e.g., cognitive abilities) based on data from the other two domains (e.g., electrophysiological and behavioral data). We iteratively predicted data from each of the three domains (electrophysiological, behavioral, and cognitive abilities data) by the other two for each new participant and each of the two models. Out-of-sample prediction was then evaluated in each of the three data domains using 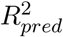 as a measure of variance explained in variables of one domain by variables from the other two domains. Note that there is no constraint of 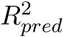 in out-of-sample evaluation to values above zero. Negative values indicate that there is more deviation of the predicted values from the true values than there is variance in the true values themselves.

#### Open-source data and analysis code

MATLAB, Python, and JAGS analysis code and data are available at https://osf.io/de75n/ and in the following repository (as of February 2018): https://github.com/mdnunez/ERPIQRT/

## Results

Mean performance (reaction times and accuracies) in the three experimental tasks is shown in **Table 1**. Grand-average waveforms of event-related potentials are presented in **Figure 4**. See **Table 2** for mean ERP latencies in both sessions.

**Table 1.** Mean RTs (SD in parentheses) for all conditions of the three experimental tasks

| Task | Session 1 |  | Session 2 |  |
| --- | --- | --- | --- | --- |
|  | Accuracies | RTs | Accuracies | RTs |
| Choice reaction time task |  |  |  |  |
| CRT2 | .99 (.01) | 382.79 (58.02) | 1.00 (.01) | 381.27 (61.01) |
| CRT4 | .99 (.01) | 477.22 (82.64) | .98 (.02) | 467.31 (85.70) |
| Recognition memory task |  |  |  |  |
| Set size 1 | .97 (.02) | 590.96 (115.67) | .98 (.02) | 584.02 (135.64) |
| Set size 3 | .97 (.02) | 728.46 (167.21) | .98 (.03) | 706.61 (176.81) |
| Set size 5 | .97 (.03) | 890.03 (240.74) | .95 (.09) | 850.98 (223.18) |
| Letter matching task |  |  |  |  |
| Physical identity | .98 (.02) | 617.79 (93.93) | .98 (.02) | 605.19 (102.41) |
| Name identity | .98 (.02) | 699.50 (113.02) | .97 (.02) | 704.38 (126.36) |

**Table 2.** Mean ERP Latencies (SD in parentheses) averaged across conditions of each of the three experimental tasks

| Task | P2 | N2 | P3 |
| --- | --- | --- | --- |
| Session 1 |  |  |  |
| Choice reaction time task | 211.54 (32.82) | 206.15 (27.71) | 330.67 (44.26) |
| Recognition memory task | 234.08 (34.48) | 251.11 (42.05) | 374.35 (74.76) |
| Letter matching task | 222.26 (33.74) | 247.87 (36.80) | 414.97 (86.45) |
| Session 2 |  |  |  |
| Choice reaction time task | 208.44 (33.77) | 210.38 (29.62) | 324.40 (42.04) |
| Recognition memory task | 230.35 (28.19) | 248.48 (43.74) | 382.39 (81.13) |
| Letter matching task | 218.16 (25.27) | 240.02 (44.65) | 377.74 (75.09) |

### In-sample prediction

The first linking model (see **Figure 5** and **Figure 6**), in which cognitive abilities were solely predicted by neural processing speed, provided an acceptable account of the training data. On average, it explained 63% of the variance in cognitive abilities tests, 62% of the variance in ERP latencies, 87% of the variance in the 25th percentile of the RT distribution, 89% of the variance in the 50th (median) percentile of the RT distribution, 83% of the variance in the 75th percentile of the RT distribution, and 30% of the variance in accuracies in reaction time tasks. Note that the cognitive latent variable model may have explained more variance in reaction times than in ERP latencies and cognitive abilities tests because the measurement model of reaction times was more complex (allowing the task-, condition- and session-specific estimation of boundary separation and non-decision time models not depicted in the structural equation model visualization) than the other two more parsimonious measurement models. The DIC of the overall hierarchical model with the first linking structure was −3.2012 * 105 and was thus the favored model by the DIC (compared to the second linking structure DIC below). The latent neural processing speed variable predicted the latent cognitive abilities variable to a large degree, *β* = .84, CI 95% [.75; .91], suggesting that participants with greater cognitive abilities showed a substantially higher neural processing speed.

The second linking model (see **Figure 7** and **Figure 5**), in which the effect of neural processing speed was partly mediated by drift rates, also provided a good account of the training data. It explained on average 63% of the variance in cognitive abilities tests, 63% of the variance in ERP latencies, 89% of the variance in the 25th percentile of the RT distribution, 90% of the variance in the 50th (median) percentile of the RT distribution, 83% of the variance in the 75th percentile of the RT distribution, and 25% of the variance in accuracies in reaction time tasks. The explained variance is therefore nearly identical to the first linking model. The DIC of the model with the second linking structure was −3.2007 * 105, a larger, and thus unfavored, DIC compared to the previous model. Again, the latent neural processing speed variable predicted the latent cognitive abilities variable, *β*_1_ = .78, CI 95% [.63; .89]. Individual latent neural processing speeds also predicted individual latent drift rates, *β*_3_ = .17, CI 95% [.05; .33]. However, there was only weak evidence that greater latent drift rates predicted greater cognitive abilities, *β*_2_ = .23, CI 95% [-.05; .52]. Nevertheless, we found some evidence for an indirect effect of neural processing speed on cognitive ability test scores that was mediated by drift rates, *β_indirect_* = .04, CI 95% [-.01; .09]. See **Figure 8** for posterior density distributions of the standardized regression weights. To compare both models, we calculated DICs as measures of model fit. The difference between DICs of ∆DIC = 43.27 indicated that the mediation model could not provide a better account of the data than the more parsimonious regression model.

**Figure 8.**
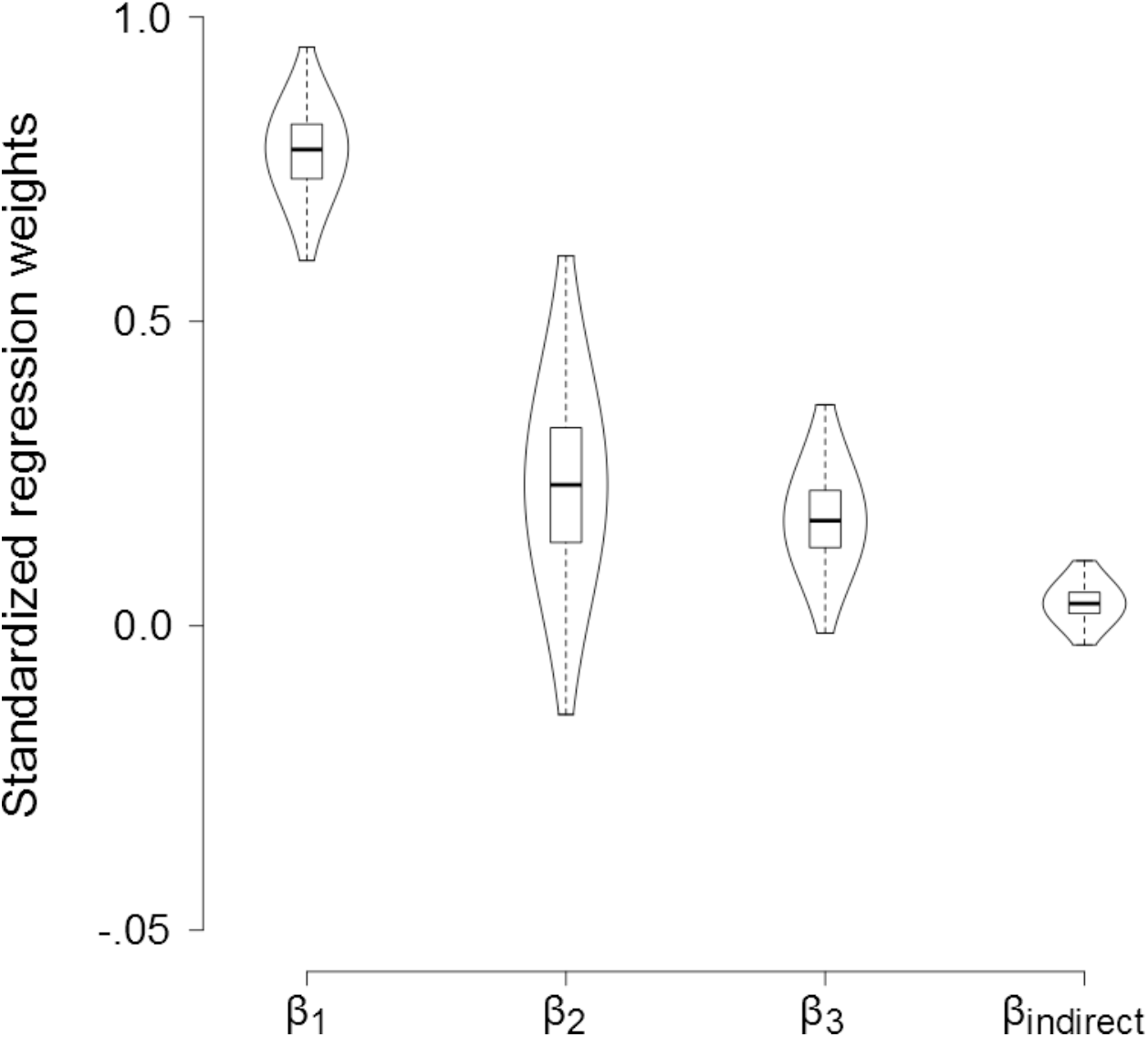
Posterior density distributions of the standardized regression weights of the mediation linking model. Boxes indicate the interquartile range with the median as a horizontal line. *β*_1_ = regression of latent cognitive abilities factor on latent neural processing speed factor; *β*_2_ = regression of latent cognitive abilities factor on latent drift rate factor; *β*_3_ = regression of latent drift rate factor on latent neural processing speed factor; *β_indirect_* = indirect effect.

### Out-of-sample prediction of new participants

To evaluate the ability to predict unknown data of a new participant in one domain (e.g., unknown cognitive ability test scores) from observed data in another domain (e.g., observed ERP latencies), we assessed out-of-sample-prediction ability for both models in a test set of 22 randomly drawn participants.

Given a *new* participant’s ERP and RT data, the regression linking model (see **Figure 6**) yielded the ability to make somewhat accurate predictions of that participant’s cognitive abilities test scores and ERP latencies. That is, out-of-sample prediction explained 39% of the variance in cognitive abilities tests across participants and tasks and 22% of the variance in ERP latencies across participants and tasks. However, out-of-sample prediction of reaction time data was not successful, *R*^2^ = *−.*51 in the 25th percentile of the RT distribution, *R*^2^ = *−.*50 in the 50th (median) percentile of the RT distribution, and *R*^2^ = *−.*67 in the 75th percentile of the RT distribution. Accuracies could also not be predicted successfully, *R*^2^ = −1.22.

The mediation linking model (see **Figure 7**) produced very similar predictions of participants’ cognitive abilities test scores and ERP latencies. Out-of-sample prediction explained 36% of the variance in cognitive abilities tests across participants and tasks and 23% of the variance in ERP latencies across participants and tasks. Again, prediction of out-of-sample reaction time data was not successful, *R*^2^ = –1.10 in the 25th percentile of the RT distribution, *R*^2^ = − .96 in the 50th (median) percentile of the RT distribution, *R*^2^ = –2.09 in the 75th percentile of the RT distribution, and *R*^2^ = –1.46 for accuracies in the reaction time tasks.

## Discussion

We investigated whether the association between neural processing speed and general cognitive abilities was mediated by the velocity of evidence accumulation. For this purpose, we used a Bayesian cognitive latent variable modeling approach that allowed the joint modeling behavioral, neural, and cognitive abilities data and estimation of relationships between higher-order latent variables. The cognitive latent variable model was able to predict a substantial amount of variance in cognitive ability test scores in new participants solely based on those participants cortical processing speeds.

We observed a strong association between neural processing speed and general cognitive abilities in the way that individuals with greater cognitive abilities showed shorter latencies of ERP components associated with higher-order cognition. Moreover, we found that individuals with greater neural processing speed also showed a greater velocity of evidence accumulation. Given an individuals speed of neural information processing and evidence accumulation, we could predict about 40 percent if their variance in intelligence test scores. However, the association between neural processing and general cognitive abilities was only mediated by drift rates to a small degree, and the more complex mediation model did not provide a better account of the data than the more parsimonious regression model.

These results support the idea that a greater speed of neural information processing facilitates evidence accumulation, and that this increase in the velocity of evidence accumulation translates to some degree to advantages in general cognitive abilities. Although previous studies reported substantial correlations between drift rates and cognitive abilities (Schmiedek et al., 2007; Schmitz & Wilhelm, 2016; van Ravenzwaaij et al., 2011), and although preliminary results suggested that measures of neural processing speed and drift rates can load onto the same factor (Schubert et al., 2015), the present study provided the first direct test of the hypothesis that the velocity of evidence accumulation mediates the relationship between neural processing speed and cognitive abilities. Our results suggest that only a small amount of the shared variance between neural processing speed and cognitive abilities can be explained by individual differences in the velocity of evidence accumulation as a mediating cognitive process. We provide three conceptual explanations why the velocity of evidence accumulation may only explain little of the natural variation in human cognitive abilities associated with cerebral processing speed.

### (1) A common latent process

Both neural processing speed and the velocity of evidence accumulation may reflect properties of the same latent process that is related to general cognitive abilities. However, the drift rate may be an impurer measure of this latent process or may be contaminated by properties of other processes unrelated to cognitive abilities. This position is supported by the observation that we found an association between ERP latencies and drift rates, and by our result that drift rates mediated the relationship between ERP latencies and cognitive abilities at least partially. Moreover, this explanation is consistent with previous research, which suggested that the P3 may be a neural correlate of the evidence accumulation process captured by drift rates (Kelly & O’Connell, 2013; O’Connell et al., 2012; Ratcliff et al., 2009, 2016; van Ravenzwaaij et al., 2017). The fact that the associations between neural processing speed and drift rates were lower than the correlations reported in the literature may be due to deviations from previous studies: First, the current study focused on ERP latencies as measures of neural processing speed, whereas previous studies analyzed the relationship between amplitude and capacity-related measures of the EEG and drift rates. Second, previous studies focused mostly on late centro-parietal potentials, whereas the current study included a more diverse time-course and topography of ERP components. Third, we only related the latent neural processing speed factor, which reflected the shared variance between different ERP latencies across different tasks, to the latent drift rate factor, and did not inspect task or component-specific correlations. Considering the psychometric properties of both ERP latencies and drift rates (Schubert et al., 2017, 2015), it is highly likely that associations between ERP latencies and drift rates would have been higher if we had modeled correlations separately for each condition of each experimental task. However, this task- or condition-specific variance in ERP latencies and drift rates is not of interest regarding general cognitive abilities.

### (2) Other candidate cognitive processes

The velocity of evidence accumulation may not be the appropriate candidate process mediating the relationship between neural processing speed and cognitive abilities. Instead, shorter latencies of ERP components associated with higher-order cognitive processing may reflect a faster inhibition of extraneous processes and may thus be a neural correlate of the efficiency of selective attention (Polich, 2007). The idea that attentional processes underlie individual differences in cognitive abilities has been discussed numerous times. Process overlap theory (Kovacs & Conway, 2016), for example, proposes that a limited number of domain-general and domain-specific cognitive processes contribute to individual differences in general cognitive abilities. In the framework of process overlap theory, attentional processes represent a central domain-general bottleneck that constrains cognitive performance across different tasks. This notion is supported by several studies reporting substantial associations between measures of attentional control and executive processes and general cognitive abilities (e.g., Unsworth, Fukuda, Awh, & Vogel, 2014; Wongupparaj, Kumari, & Morris, 2015).

Additionally, a greater neural processing speed may directly facilitate the storage and updating of information in working memory (Polich, 2007), and may thus lead to a greater working memory capacity, which may positively affect performance in a large number of cognitive tasks. This notion is supported by numerous studies reporting large and even near-unity correlations between measures of cognitive abilities and working memory capacity (e.g., Conway, Cowan, Bunting, Therriault, & Minkoff, 2002; Engle, Tuholski, Laughlin, & Conway, 1999; Kyllonen & Christal, 1990). Individual differences in these working memory processes may not be reflected in drift rates estimated in simple binary decision tasks. Instead, future studies could use mathematical models of working memory, such as mathematical implementations of the time-based resource sharing model (Barrouillet, Bernardin, & Camos, 2004) or the SOB-CS (Oberauer, Lewandowsky, Farrell, Jarrold, & Greaves, 2012), to explicitly model individual differences in parameters of working memory processes and relate these parameters to neural data in a cognitive latent variable model.

Finally, it might even be possible that several cognitive processes mediate the relationship between neural processing speed and cognitive abilities, and that parameters of each single cognitive process only account for a small amount of the substantial association. Larger multivariate studies incorporating cognitive models of these candidate cognitive processes would be required to quantify additive and multiplicative effects of different cognitive processes on the relationship between neural processing speed and general cognitive abilities.

### (3) Brain properties as confounding variables

Individual differences in neural processing speed may reflect structural properties of the brain that give rise to individual differences in cognitive abilities. Brain properties may be related both to neural processing speed and general cognitive abilities and may thus explain the substantial association between the two variables. Previous research has shown that individuals with greater cognitive abilities showed greater nodal efficiency in the right anterior insula and the dorsal anterior cingulate cortex (Hilger, Ekman, Fiebach, & Basten, 2017). These brain regions are core components of the salience network that is assumed to be responsible for the detection of salient information and its evaluation with regard to behavioral relevance and an individuals goals (Downar, Crawley, Mikulis, & Davis, 2002; Menon & Uddin, 2010; Seeley et al., 2007). Dynamic source imaging and lesion studies have revealed that the relative timing of responses of the anterior insula and the dorsal anterior cingulate cortex to stimuli can be indexed by the N2b/P3a component of the ERP, followed by an elicitation of the P3b in neocortical regions in response to the attentional shift (Menon & Uddin, 2010; Soltani & Knight, 2000). Hence, a more efficient functional organization of the salience network may affect the timing of these ERP components and may also positively affect performance in cognitive ability tests by facilitating the goal-driven selection of task-relevant information.

#### Cognitive latent variable models

The use of cognitive latent variable models allows the simultaneous modeling of cognitive, neural and behavioral data across different tasks and ability tests. CLVMs thus allow estimating latent correlations between different measurement areas that are free of unsystematic measurement error. This property is particularly useful when dealing with time-related electrophysiological data, which have been shown to be very inconsistent in their reliability (Cassidy, Robertson, & O’Connell, 2012; Schubert et al., 2017). Moreover, CLVMs allow modeling the shared variance between diffusion model parameters across different tasks and conditions in a hierarchical way and can thus solve the problem of low-to-moderate consistencies of model parameters in individual differences research (Schubert et al., 2016).

Three advantages of the hierarchical Bayesian approach have been highlighted by the present study: First, the CLVM demonstrated advantages over classical structural equation modeling approaches in its predictive abilities in small-to-moderate sample sizes. The model has been developed based on only 92 participants and has successfully predicted 62 to 89 percent of the within-sample variance in neural, behavioral and cognitive abilities data. A conventional structural equation model with the same number of free parameters would require a substantially larger sample size. Following the rule of thumb to collect at least five observations per estimated parameter (Bentler & Chou, 1987), the same model would require a sample size of at least 480 participants in a conventional SEM framework. Taking into account the ratio of indicators to free parameters *r* (*r* = number of indicators/number of free parameters), a sample size of at least 930 participants would be required according to the equation *n* = 50 · *r*^2^ − 450 · *r* + 1100 proposed by Westland (2010) based on the simulation results by Marsh, Hau, Balla, and Grayson (1998). Such large sample sizes are hardly feasible for neuroimaging research except in large-scale collaborative research projects. The Bayesian approach presented here enabled us to fit a structural equation model of great complexity to a sample of only 92 participants. Most importantly, one of the main results previously shown in a more parsimonious conventional structural equation model applied to the same data set (i.e., the great association between neural processing speed and cognitive abilities reported by Schubert et al., 2017), was adequately recovered by the Bayesian model. Therefore, hierarchical Bayesian models may provide the neuroimaging community with a means to benefit from the advantages of structural equation modeling or other multivariate regression approaches.

Moreover, the latent drift rate trait and task-, condition-, and state-specific boundary separation and non-decision time parameters could account for nearly 90 percent of the in-sample reaction time data. In comparison, latent diffusion model parameter traits have been shown to account for only 30 to 79 percent of variance in single-task parameter estimates in a conventional structural equation model (Schubert et al., 2016). This in-sample prediction ability demonstrates that it may be beneficial to model only parameters with known trait properties (e.g., drift rate, see Schubert et al., 2016) as hierarchical factors, while the other model parameters that are known to be more strongly affected by task-specific influences (e.g., non-decision time and boundary separation, see Schubert et al., 2016) are estimated separately for each task and condition.

Second, both the cognitive model and the structural model were fitted to the data in a single step, allowing an accurate representation of parameter uncertainty in posterior distributions (Vandekerckhove, 2014), whereas previous studies relating diffusion model parameters to cognitive abilities tests have relied on a two-step process (e.g., Schmiedek et al., 2007; Schmitz & Wilhelm, 2016; Schubert et al., 2015).

Third, posterior distributions of model parameters were used to predict cognitive ability test scores from neural and behavioral data in a second independent sample. This is the first study to show that posterior predictives of regression weights relating ERP latencies, behavioral data, and cognitive ability test scores may be used to successfully generalize predictions to another independent sample and to predict a substantial amount of new individuals cognitive ability test scores solely based on their electrophysiological and behavioral data.

The model developed in the present study can be easily adjusted to include different sources of neural data, such as functional magnetic resonance imaging or diffusion tensor imaging data, and to relate these data to diffusion model parameters and cognitive ability tests. Within the same hierarchical framework, parameters of different cognitive models could be related to neural and cognitive abilities data. This would, for example, allow testing hypotheses about the relationship between parameters of working memory processes and neural and cognitive abilities data. The flexibility of the hierarchical Bayesian approach allows specifying model and linking structures directly guided by theoretical assumptions, which in turn allows direct comparisons of contradicting theories.

#### Limitations

One limitation of the present study is that the tasks used to assess individual differences in the efficiency of information processing are so-called elementary cognitive tasks. Elementary cognitive tasks are cognitively relatively undemanding tasks typically used in individual differences research to minimize the influence of individual differences in strategy use and of previous experience with these tasks on task performance. However, cognitively more demanding tasks might yield a stronger association between the velocity of evidence accumulation and cognitive abilities. Whether drift rates based on performance in more demanding tasks such as working memory tasks mediate the association between neural processing speed and cognitive abilities, remains an open question. In addition, low error rates may have limited the estimation and interpretation of diffusion model parameters. In particular, identifying drift rate and boundary separation parameters becomes difficult in tasks with few incorrect responses. Although diffusion model parameters provided a good account of the behavioral data in all three tasks, drift rate parameters might have reflected participants *decision times* to a larger degree than their *evidence accumulation rates*.

#### Conclusion

We used a cognitive latent variable model approach to show that a higher neural information processing speed predicted both the velocity of evidence acquisition and general cognitive abilities, and that a small part of the association between neural processing speed and cognitive abilities was mediated by individual differences in the velocity of evidence accumulation. The model demonstrated impressive forecasting abilities by predicting 35 to 40 percent of the variance of individual cognitive ability test scores in an entirely new sample solely based on their electrophysiological and behavioral data. Our results illustrate, however, that the assumption of a unidirectional causal cascade model, in which a higher neural processing speed facilitates evidence accumulation, which may in turn give rise to advantages in general cognitive abilities, was not supported by the data. Instead, our results support the view that neural correlates of higher-order information-processing and drift rates reflect the same latent process or structural brain property (Kelly & O’Connell, 2013; O’Connell et al., 2012; Ratcliff et al., 2009, 2016; van Ravenzwaaij et al., 2017), which may be strongly related to cognitive abilities, but that measures of neural processing speed are better measures of this property or process than drift rates. Moreover, our results suggest that individual differences in the speed of neural processing might affect a plethora of higher-order cognitive processes, that might only in concert explain the large association between neural processing speed and cognitive abilities.

## Acknowledgments

The authors thank Gidon T. Frischkorn for helpful comments on an earlier draft of this manuscript. Ramesh Srinivasan and other members of the Human Neuroscience Laboratory are well appreciated for their constructive criticism on initial work related to this manuscript.

## Funding

ALS was supported by the G.A.-Lienert-Foundation. MDN and JV were supported by NSF grant #1658303.

## Conflict of interest statement

The authors declare that the research was conducted in the absence of any commercial or financial relationships that could be construed as a potential conflict of interest.

